# Autism Polygenic Score Is Associated With Sex-Dependent Broadening of Brain Network Variability

**DOI:** 10.64898/2026.08.18.745469

**Authors:** Joe Bathelt, Danae Mitsea, Hilde M. Geurts

**Affiliations:** Dutch Autism & ADHD Research Centre, University of Amsterdam, Nieuwe Achtergracht 129-B, Amsterdam, 1018 WS, North Holland, The Netherlands; Department of Psychiatry, Amsterdam University Medical Centre, De Boelelaan 1117, Amsterdam, 1081 HV, North Holland, The Netherlands

**Keywords:** Autism, Polygenic Score, Brain Connectivity, Imaging Genetics, Social Cognition

## Abstract

**Background:** Autism polygenic scores (PGS) reliably predict case–control status yet explain little variance in autism-related traits. Landscape accounts of neurodevelopmental diversity propose that genetic liability broadens the range of viable neural configurations rather than shifting brain organisation toward dysfunction. We tested whether autism polygenic load is associated with increased variability in functional network organisation among non-autistic adults.

**Methods:** We analysed resting-state functional connectivity from 910 non-autistic adults (aged 22–35) in the Human Connectome Project. Polygenic scores were derived from the iPSYCH autism GWAS at a pre-specified threshold (***p* = 0.1**). Modularity (segregation) and global efficiency (integration) were computed at a pre-selected parcellation size and density (100-node, 20%), and residualised for age, intracranial volume, and head motion. Variance effects were assessed by variance regression including a PGS × sex interaction, decile-stratified dispersion trends, and PGS-balanced bootstrap resampling. Edge-wise analyses used false discovery rate correction.

**Results:** Modularity variability broadened with polygenic load in a sex-dependent manner (sex × PGS ***β* = 1.92 × 10*^−^*^4^**, ***p* = 0.031**). Decile trends (**M − F** difference **= 0.82**, ***p* = 0.034**) and balanced-bootstrap trends (difference **= 1.19**, ***p* = 0.032**) both differed by sex: variance increased across polygenic bins in males (***r* = 0.57**, one-tailed ***p* = 0.021**) but not females. No comparable effect emerged for global efficiency (all ***p* ≥ 0.54**). Polygenic scores showed no association with social-cognitive difficulty (***β* = 0.11**, ***p* = 0.209**), mean network organisation, or connectivity after correction.

**Limitations:** All participants were non-autistic adults and the analysis was cross-sectional. The identified effects are small and the sample size not sufficient to resolve very small effects often reported in genetics studies. Characterisation of genetic effects in women may be influenced by biases in the data used to calculate polygenic scores.

**Conclusions:** Autism polygenic load broadened modular network configurations in males without shifting mean organisation or its behavioural correlates, offering partial support for landscape accounts.

## 1 Background

Genetic predisposition to neurodevelopmental conditions presents a fundamental puzzle: variants associated with clinical diagnoses are continuously distributed across the general population[1], yet most individuals carrying elevated genetic burden never develop impairing symptoms. Even among individuals in the top decile of autism polygenic scores, the prevalence of autism diagnosis increases only 2.3-fold compared to population base rates[2]. Further, autism polygenic scores explain less than 5% of variance in associated behavioural traits[1]. This discontinuity between genetic risk and phenotypic outcome suggests that the pathway from genome to behaviour is not deterministic, but the mechanisms underlying this variability remain poorly understood. One possibility is that genetic predisposition uniformly shifts brain organisation toward dysfunction, with clinical outcomes emerging only when deviation exceeds some threshold. An alternative view, grounded in recent network neuroscience theory, proposes that genetic predisposition instead expands the range of possible neural configurations, thereby increasing variability rather than systematically altering brain organisation.

The connectome landscape theory of dysconnectivity provides a formal framework for this latter perspective[3]. Rather than conceptualising brain conditions as patterns of aberrant connectivity, this theory proposes that clinical groups occupy a broader region of the possible “connectome landscape” defined by the balance between network integration and segregation. Building on this framework, Finn and colleagues articulated an Anna Karenina principle for brain organisation: whereas optimal network configurations may be constrained to a narrow region of this landscape, suboptimal configurations can deviate in multiple directions[4]. Under this view, genetic predisposition does not push brain development toward a particular dysfunction, but rather loosens the constraints that typically channel neural organisation toward a limited set of optimal solutions.

This landscape perspective generates distinct predictions that can be empirically tested. If genetic predisposition shifts brain organisation toward dysfunction, individuals with higher polygenic load should exhibit systematic differences in mean connectivity patterns and associated behavioural difficulties. In contrast, if genetic predisposition expands developmental possibilities, we should observe increased variability in neural organisation without corresponding shifts in group means. Further, behavioural outcomes should depend not on the genetic load directly, but on whether individual developmental trajectories achieve optimal configurations within the expanded landscape. Critically, these predictions apply not only to individuals with clinical diagnoses, but across the continuum of genetic predisposition present in the general population.

Autism spectrum conditions offer an ideal test case for adjudicating between these competing models. Large-scale genome-wide association studies have identified hundreds of common genetic variants associated with autism, enabling construction of polygenic scores that quantify cumulative genetic predisposition[5]. These scores robustly predict case-control status, yet explain only modest variance in behavioural outcomes within either clinical or population samples[6]. This limited predictive capacity is precisely what landscape theory would anticipate: if genetic predisposition expands variability rather than shifting means, aggregate scores should poorly predict individual outcomes despite capturing genuine biological signal.

Emerging neuroimaging evidence supports the prediction that autism is associated with increased neural idiosyncrasy. Analysing functional connectivity data from the Autism Brain Imaging Data Exchange (ABIDE)[7], Benkarim and colleagues found that autistic individuals showed significantly increased idiosyncrasy in default mode, somatomotor, and attention networks compared to non-autistic controls[8]. This functional idiosyncrasy correlated with symptom severity and co-localised with expression patterns of autism risk genes, suggesting a direct link between genetic factors and neural variability. In the same data, Dickie and colleagues demonstrated greater variability in the spatial locations of resting-state networks in autism, with non-autistic individuals showing a developmental decrease in variability from childhood to adulthood that was absent in autistic individuals[9]. Xie and colleagues, also using ABIDE, found significantly lower person-based similarity in grey matter volume profiles among autistic compared to non-autistic individuals[10]. Importantly, these patterns replicate in independent samples. In the EU-AIMS Longitudinal European Autism Project, Ecker and colleagues demonstrated that autistic individuals’ total degree of neuroanatomical atypicality correlated with polygenic scores for autism and scaled with symptom severity[11]. Using a study-specific Taiwanese sample, Tung and colleagues found significant white matter tract idiosyncrasy in autism that was notably absent in non-autistic siblings, suggesting that increased variability reflects more than familial background[12]. Together, these findings spanning functional connectivity, cortical morphometry, and white matter organisation across multiple independent samples document the increased neural variability that landscape theory predicts when genetic predisposition expands the developmental landscape.

However, interpreting variability findings in clinical samples remains complicated by potential confounds. Greater neural diversity in diagnosed individuals could reflect genuine biological variation, but might also arise from heterogeneous clinical presentations[13], varying comorbidities[14], or different treatment histories[15]. A further methodological concern is that much of the existing evidence derives from retrospective data collections such as ABIDE, which aggregated neuroimaging data across sites without prior harmonisation of acquisition protocols or clinical characterisation[7]. Site-related variance in such collections may inflate apparent neural idiosyncrasy, making it difficult to distinguish protocol heterogeneity from genuine biological variability. Disentangling meaningful neural variation from these confounds requires examining genetic predisposition effects in prospectively collected samples with standardised protocols. The Human Connectome Project[16] provides an unprecedented opportunity to test landscape theory predictions at scale, offering comprehensive genetic, neuroimaging, and behavioural data from over 900 adults without psychiatric or neurological diagnoses, all acquired using identical scanning parameters and quality control procedures.

The present study leverages this resource to directly test competing predictions about how autism genetic predisposition influences brain organisation. We computed polygenic scores from the largest available autism genome-wide association study[5] and examined their relationship with functional brain network organisation, assessed through both edge-wise connectivity analyses and graph-theoretic measures of network integration (global efficiency) and segregation (modularity). Our central prediction, derived from landscape theory, is that individuals with higher polygenic scores will exhibit greater variability in brain network organisation, specifically in measures indexing the integration-segregation balance, without corresponding shifts in mean organisation.

## 2 Methods

### 2.1 Participants

The analysis drew upon data from the Human Connectome Project Young Adult (HCP-YA) 1200 Subjects release[16], a large-scale initiative that collected comprehensive behavioural, genetic, and neuroimaging data from 1,206 healthy adults between 22 and 35 years of age. Participants were recruited between 2010 and 2016 in the United States, primarily from Missouri with additional recruitment from surrounding states to ensure diverse representation in terms of race, ethnicity, and socioeconomic status. The recruitment was designed to reflect the demographic makeup of the U.S. population as represented in the 2000 decennial census[16]. The inclusion criteria required participants to be generally healthy and without significant medical or psychiatric conditions, including neurodevelopmental disorders or chronic diseases. Some participants were excluded at different steps because of missing data or failed data quality control (see Figure S1 for a data retention flowchart). The final sample consisted of 910 participants.

### 2.2 Social processing factor

To create a composite behavioural outcome measure for our main analysis, we developed a social processing factor score from existing HCP-YA measures to capture autism-relevant variation. We selected numeric measures related to social, emotional, or language processing from task assessments and NIH Toolbox questionnaires, including: Penn Emotion Recognition 40 (facial emotion recognition), Language processing (reasoning about character intentions in Aesop’s fables), Emotion processing (matching facial expressions), Social processing (interpreting meaningful vs. random shape interactions), and NIH Toolbox Social Relationships questionnaire (6 domains of social functioning; see Supplementary Materials for detailed descriptions). Data preprocessing involved, scaling variables, marking outliers as missing (outside 1.5×IQR, Penn Emotion Recognition 40: n=30 [2.93%]; Language processing: n=58 [5.66%]; Emotion processing: n=59 [5.76%]; Social processing: n=71 [6.93%]), and imputing missing values using gradient boosting models (miceforest v.5.6.2, doi:10.5281/zenodo.7428632). The sample was split into training (80%) and test (20%) sets. Exploratory factor analysis was conducted using the `psych` package; confirmatory factor analysis was carried out using `lavaan` v.0.6.15.

We tested five different combinations of behavioural measures using exploratory factor analysis (EFA) on training data, followed by confirmatory factor analysis (CFA) validation on test data (see Table 2.2). The reaction time (RT) Model, which included RT measures from the four social-cognitive tasks, was selected based on superior fit indices and theoretical interpretability. While the Relative RT Model (RTs adjusted by subtracting 0-back task RT) showed marginally better fit statistics, the adjustment reduced between-task variability that may be theoretically important for capturing individual differences in social-cognitive processing. CFA on held-out data (*n* = 205) confirmed excellent model fit with all factor loadings significant (*p <* 0.01) and standardized loadings ranging from 0.30 to 0.70 (Penn Emotion Recognition 40: 0.30; Language processing: 0.34; Emotion processing: 0.44; Social processing: 0.70). The factor explained an average of 22% of variance across indicators, with the highest loading on the social and emotion processing tasks (Penn Emotion Recognition 40: 0.09; Language processing: 0.11; Emotion processing: 0.20; Social processing: 0.49). Based on these loadings and theoretical interpretation of the tasks, we propose that the factor score captures social processing, with higher factor scores indicating greater social-cognitive difficulties.

**Table 1.** Model Fit Statistics for Confirmatory Factor Analysis.

| Model | RMSEA | TLI | $p$-value | BIC |
| --- | --- | --- | --- | --- |
| RT Model | 0.008 | 0.998 | 0.347 | −11.3 |
| Relative RT Model | 0.000 | 1.000 | 0.462 | −11.9 |
| Questionnaire Model | 0.167 | 0.820 | <0.001 | 154.0 |
| RT + Questionnaire Model | 0.110 | 0.783 | <0.001 | 148.0 |
| Relative RT + Questionnaire Model | 0.120 | 0.748 | <0.001 | 215.0 |

The social difficulties score showed small to moderate correlations with other composite scores. The strongest associations were with fluid intelligence (*r* = −0.34*, p* = 9.38e−28) and processing speed (*r* = −0.31*, p* = 5.01e−22), followed by card sorting (*r* = −0.29*, p* = 3.40e−19). Weaker but significant correlations were observed with picture vocabulary (*r* = −0.09*, p* = 0.007) and list-sorting working memory (*r* = −0.10*, p* = 0.003; all with *n* = 948). The negative correlations indicate that higher social difficulty scores (reflecting longer reaction times and greater social difficulties) are associated with lower accuracy scores on general cognitive measures. The small-to-moderate correlations suggest that the social difficulties score captures unique variance, putatively related to social processing, which is not captured by an overarching cognitive composite scores.

### 2.3 Polygenic Score (PGS) Calculation

Base data were obtained from the iPSYCH autism genome-wide association study [5], in which autism PGS explained 2.45% of case-control variance in the original discovery sample. Base data quality control excluded duplicate variants and retained SNPs with minor allele frequency (MAF) ≥ 0.01 and imputation quality score ≥ 0.7. Target genotype data from the Human Connectome Project (HCP) comprised 1,134 individuals with 472,278 SNPs on genome build GRCh37. Target data underwent quality control using the `plinkQC` package [17], with thresholds applied at the individual level (sample missingness, sex-chromosome inconsistencies, heterozygosity outliers) and at the marker level (SNP missingness, Hardy–Weinberg equilibrium, MAF); full parameter values are reported in Supplementary Methods S2 and Table S1. Linkage disequilibrium (LD) pruning was performed with a 100-SNP window, 25-SNP step, *r*^2^ > 0.5. Strand alignment and allele recoding against the base data were performed prior to scoring (Supplementary Methods S3).

Due to the risk of overfitting with limited sample sizes, we pre-specified the threshold at *p* = 0.1, the optimal threshold reported in the discovery GWAS [5] rather than optimising the PGS threshold against our target outcome. Using the discovery-GWAS threshold is the most principled option available, because it reflects the polygenic architecture of autism as estimated in the largest clinical sample to date and uses no information from our own target data. To calibrate PGS in an unrelated subset before extension to the full sample, we computed scores using PRSice v2.3.5 [18] in participants passing both phenotypic non-twin status and an IBD-based relatedness cutoff of *π̂* < 0.125, restricted to those with complete social processing factor scores (*n* = 160; see Supplementary Methods S5). Five-fold cross-validated correlation between PC-corrected PGS and the social processing factor in this calibration subset was *r* = 0.04 (*SD* = 0.11).

To extend PGS estimates to the full dataset including related individuals, we implemented a Best Linear Unbiased Prediction (BLUP) approach using GCTA v1.94.3 [19]. A genomic relationship matrix was computed for the full quality-controlled sample on autosomal markers; a linear mixed model was fitted with the unrelated-sample PGS as the phenotype to obtain random-effect predictions for all individuals; per-SNP BLUP effect sizes were then back-calculated and applied to the full sample using PLINK’s weighted scoring [20] to yield the final BLUP-extended PGS (Supplementary Methods S6). The BLUP-extended PGS and the original PRSice-derived PGS were in very high agreement within the unrelated subset (*r* = 0.99, *p* = 4.47 × 10*^−^*^126^; Figure S2), with comparable distributions. For downstream association analyses, the BLUP-extended PGS was residualised for age and the first 5 principal components, and the residuals were standardised.

### 2.4 Processing of functional connectivity data

We utilised functional connectivity matrices provided in the HCP s1200 release. The full details of the preprocessing are described in the HCP documentation [21]. In short, resting-state fMRI data from 1003 participants with 4 complete runs were processed. After removing participants with missing confound data or high motion (relative RMS > 0.2), and further restricting to individuals with complete behavioural and polygenic score data, the final analytical sample comprised 910 participants (see Figure S1 for a flowchart of data retention).

Each participant’s 15-minute resting-state runs underwent custom minimal preprocessing [22] and artefact removal using ICA+FIX. The data were temporally demeaned with variance normalization. Group-level parcellation was performed using FSL’s MELODIC tool. First, MIGP (MELODIC’s Incremental Group-PCA) generated the top 4500 weighted spatial eigenvectors from group-averaged PCA [21]. This output was fed into spatial Independent Component Analysis (ICA) at multiple dimensionalities (15, 25, 50, 100, 200, and 300 components), applied in grayordinate space. The resulting ICA maps functioned as data-driven parcellations. For each participant, node time series were extracted using the dual-regression approach [23], where ICA spatial maps served as regressors against individual subject data to estimate one representative time series per network node. Network connectivity matrices (”netmats”) were created using the FSLNets toolbox with Pearson correlations between node time series.

### 2.5 Graph Theory Analysis

To operationalise the integration and segregation dimensions of the landscape theory [3], we computed modularity (segregation) and global efficiency (integration) at multiple network densities [24]. Individual connectivity matrices were thresholded proportionally to retain the top 20% of strongest connections for primary analyses, with sensitivity analyses conducted at 15% and 25% network densities to assess robustness across sparsity levels. These thresholds are higher than typically reported range of 2-10% for resting-state fMRI studies [25]. However, our own analysis showed that networks fractionated at thresholds below 10%, which is likely related to the independent component decomposition used to calculate the HCP netmats, as ICA-derived networks tend to be sparser [26].

Community structure was defined using consensus modularity clustering applied to the group-average connectivity matrix. We implemented signed modularity detection using the Louvain algorithm from the Brain Connectivity Toolbox (Python implementation v0.6.1), running 50 iterations with adaptive resolution parameter selection (*γ* = 1.0 − 3.0) to optimize community detection. The consensus partition was determined using a majority-rule threshold (*τ* = 0.5), and the final community assignment showing the highest modularity was retained.

The parcellation resolution used for all subsequent network analyses was selected *a priori* on the basis of principled criteria applied to the consensus community-detection results, thereby avoiding the circularity that arises when resolution is chosen by the presence of effects of interest. We evaluated all available ICA-derived parcellations (15, 25, 50, 100, 200, and 300 components) against two pre-specified criteria: (1) the number of detected communities should fall within 5–15, consistent with the canonical number of intrinsic resting-state networks reported in prior work [27]; and (2) among parcellations meeting this community-count target, we selected the solution with the highest joint stability–modularity score (defined as 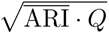, where ARI is the mean adjusted Rand index across 50 consensus iterations and *Q* is the signed modularity of the consensus partition). Only the 100-node parcellation met the community-count target (6 communities; 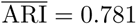; *Q* = 0.458; joint score = 0.598) and was therefore designated as the primary parcellation for all network analyses reported here. The resulting community solution showed good separation and aligned with the expected large-scale network organisation (see Figure 1).

**Fig. 1.**
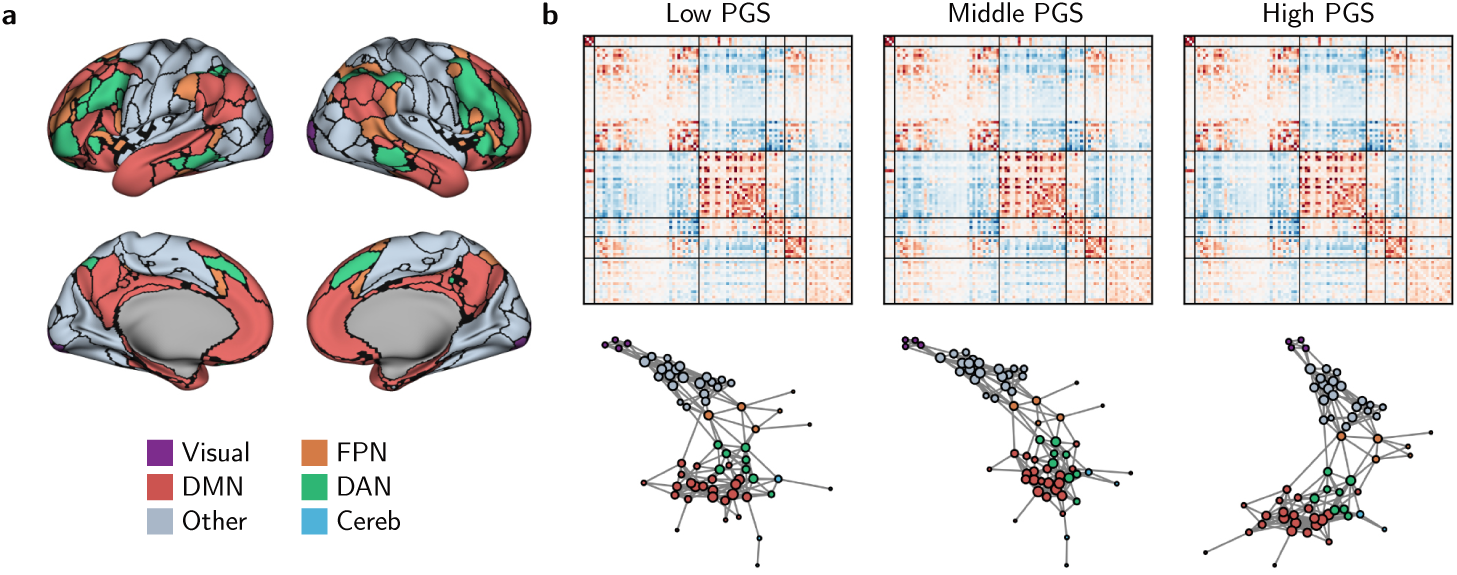
Brain network organisation by polygenic score (PGS) group. (a) Cortical and subcortical parcellation showing the anatomical organisation of brain regions used for network analysis based on consensus community detection applied to the group-average network. Different colours represent distinct functional networks. (b) Averaged connectivity matrices (top) and spring-embedded network visualizations (bottom) for low, middle, and high PGS groups at 5% network density. Matrices and networks represent the mean across 1,000 bootstrap samples to control for group size differences. Abbreviations: Visual – Visual network; DMN – Default Mode Network; FPN – Fronto-Parietal Network; DAN – Dorsal Attention Network; Cereb – Cerebellar Network.

This group-level community structure was then applied to calculate individual modularity scores using signed modularity measures that preserve both positive and negative correlations, representing how well each participant’s connectivity pattern conformed to the group-average network organisation. We opted to use signed modularity to preserve both positive and negative correlations, as altered anti-correlations between canonical networks (e.g., default mode and task-positive networks) are a key feature of connectivity differences in autism [28–30]. See Figure 1 for a visualisation of the community solution.

Global efficiency was calculated as the average inverse shortest path length between all node pairs, providing a measure of network integration and information transfer capacity [31]. Following standard practice, negative correlations were set to zero prior to path length calculations, as negative weights can produce undefined shortest paths in efficiency computations [24]. Connection weights were then converted to connection lengths.

We assessed continuous heteroscedasticity using four convergent approaches. The Breusch-Pagan and White tests examined whether residual variance from a linear regression of the network measure on PGS depended systematically on PGS, with the latter additionally accounting for non-linear dependencies. Quantile regression estimated the relationship between PGS and the network measure at multiple quantiles of the outcome distribution (0.10, 0.25, 0.50, 0.75, 0.90), with significantly diverging slopes across quantiles indicating fan-shaped heteroscedasticity consistent with landscape-theory predictions. Finally, PGS-balanced bootstrap resampling (1,000 iterations) partitioned the sample into five PGS quintile bins and resampled equal-sized subsamples within each, eliminating differences in observation density across the spectrum that could otherwise inflate apparent variance trends; we then tested whether the bootstrap-derived correlation between mean PGS per bin and within-bin variance significantly exceeded zero (one-tailed test).

At the pre-specified 100-node parcellation, we conducted sensitivity analyses to assess robustness across network density thresholds (15%, 20%, and 25%). We quantified consistency as the percentage of thresholds showing significant effects (*p <* 0.05) and established a robustness criterion of ≥ 80% consistency for core findings.

Prior to all variance, heteroscedasticity, and brain-behaviour analyses, modularity and global efficiency values were residualised against age, intracranial volume (Freesurfer-derived), and head motion (mean frame-wise displacement) using ordinary least squares regression; the resulting residuals were used as the network metrics in all subsequent tests. Covariate adjustment in the edgewise connectivity analysis is described separately below.

### 2.6 Edgewise Analysis

Connectivity features were defined by flattening the upper triangle of each correlation matrix. Each feature was robust-scaled, then residualised by regressing out covariates (age, sex, intracranial volume, head motion) using linear models. Social difficulty scores and polygenic scores were z-scored. An interaction term was computed for each connectivity feature by multiplying its residualised value with the standardised polygenic score, creating a set of interaction predictors.

We conducted three primary univariate analyses: (1) correlations between residualized connectivity features and social difficulty scores to identify brain networks associated with social functioning, (2) correlations between residualized connectivity features and polygenic risk scores to examine the neural correlates of genetic predisposition, and (3) correlations between interaction terms (connectivity × polygenic score) and social difficulty scores to test for gene-brain-behaviour interactions. For each analysis, we computed Pearson correlation coefficients across all pairwise connectivity features derived from the upper triangle of the connectivity matrix. For group comparisons, we calculated t-statistics and effect sizes (Cohen’s d) to identify connections that differed significantly between groups. Multiple comparisons correction across connectivity features was applied using the false discovery rate (FDR) method of Benjamini-Hochberg with an alpha level of 0.05.

### 2.7 Use of AI Tools

Portions of the analysis code in this study were developed with the assistance of Claude Code (Anthropic), using Claude Opus 4.8 as the underlying model, between February and July 2026. The tool was used to scaffold, refactor, and debug scripts in the phenotypic preprocessing, genetics, and connectivity pipelines. All AI-assisted code was reviewed, tested, and validated by the lead author (JB), who takes full responsibility for its correctness. No AI tool was used to generate the scientific interpretation or manuscript text beyond standard copy-editing assistance.

## 3 Results

### 3.1 Genetic Predisposition Is Associated with Increased Brain Network Variability

Landscape theory predicts that genetic predisposition expands the variability of viable neural configurations without systematically shifting their central tendency. To evaluate this prediction at the network level, we examined two complementary graph-theoretic properties were analaysed: modularity, which indexes network segregation, and global efficiency, which indexes network integration. Both were evaluated at a pre-specified 100-node parcellation and 20% density threshold. Network metrics were residualised for age, intracranial volume, and head motion prior to analysis. An initial variance model that also included sex as nuisance regressor yielded an apparent positive association between polygenic load and modularity variability. However, subsequent quality control review traced this effect to incorrect chromosomal sex assignment in the HCP genotype data, which had caused all but one female participant to be removed by the upstream sex-consistency filter; once the filter was corrected and both sexes were retained, the unconditional polygenic score (PGS) effect no longer held. All variance models reported below therefore include sex as a main effect and a PGS × sex interaction.

Across the full analytic sample (*N* = 910), three complementary tests converged on a sex-dependent broadening of modularity variability with increasing polygenic load. Variance regression of squared modularity residuals on PGS, sex, and their interaction revealed a significant sex × PGS interaction (*β* = 1.92×10*^−^*^4^, *p* = 0.031), even though the omnibus heteroscedasticity test did not reach significance (LM = 5.09, *χ*^2^(3), *p* = 0.166). Decile-stratified standard-deviation trends differed significantly between sexes (M − F trend difference = 0.82, bootstrap *p* = 0.034), with males exhibiting a positive association between polygenic decile and modularity variability (*r* = 0.46) and females exhibiting a non-significant negative trend (*r* = −0.36). PGS-balanced bootstrap analyses, which control for any imbalance in observation density across the genetic spectrum, demonstrated a similarly sex-specific pattern (M − F trend difference = 1.19, *p* = 0.032): in males, variance increased reliably across polygenic bins (trend *r* = 0.57, one-tailed *p* = 0.021), whereas in females no such expansion was observed (trend *r* = −0.62, n.s.). Figure 2 presents this broadening within the male subsample (*n* = 429), in whom the effect is concentrated: the dispersion of modularity widens across the low, middle, and high PGS reference groups, whereas global-efficiency dispersion does not.

**Fig. 2.**
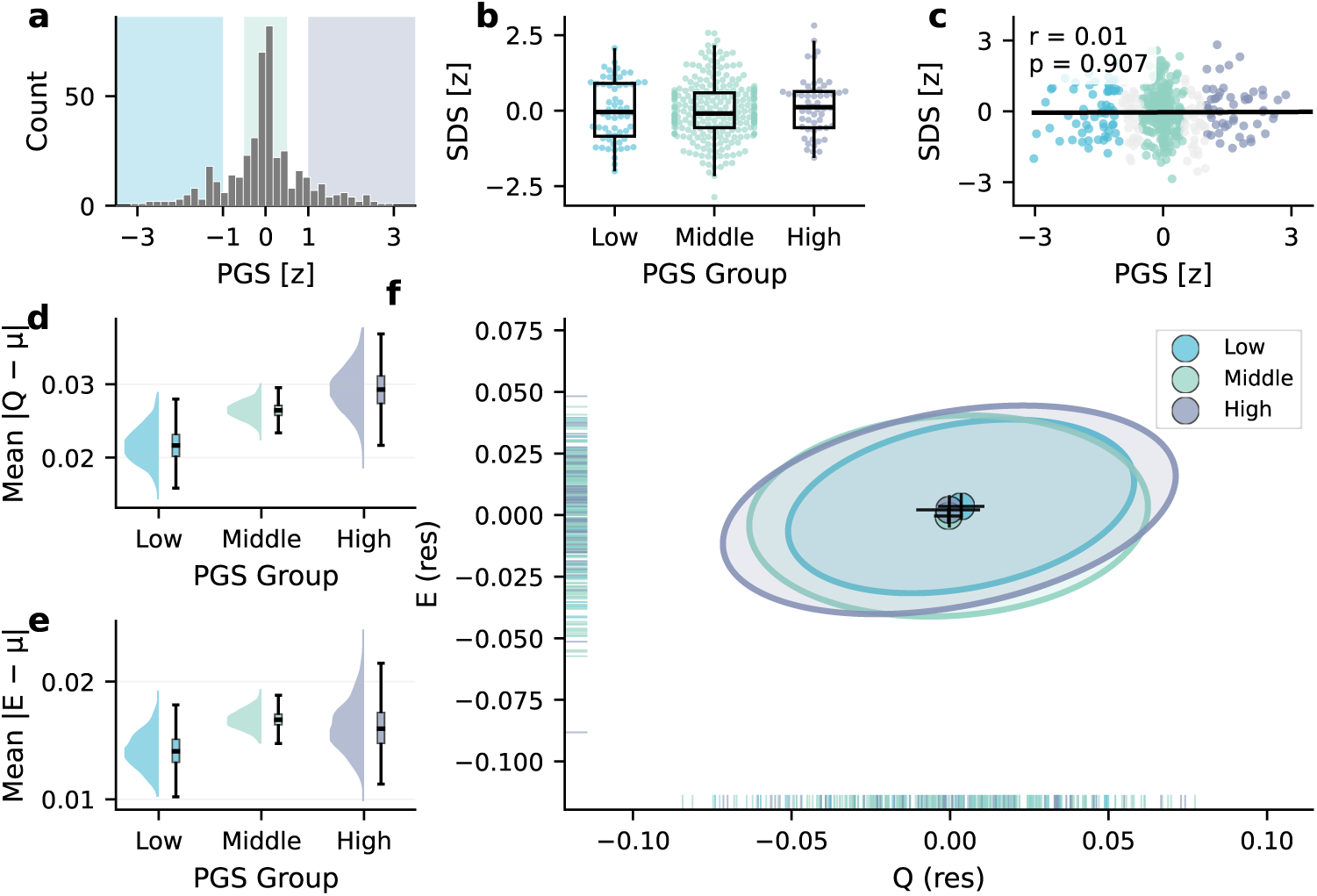
Broadening of modularity variability with increasing autism polygenic load in the male subsample. All panels show the male subsample (n = 429), in whom the sex-dependent broadening of modularity variability is concentrated (see main text and Discussion). (a) Distribution of standardised autism polygenic scores (PGS), with the *low* (*z* < −1), *middle* (*−*0.5 *≤ z ≤* 0.5), and *high* (z > 1) reference groups shaded. (b) Social difficulty score (SDS) by PGS reference group. (c) Association between PGS and SDS, shown as individual participants with the ordinary-least-squares regression line and Pearson correlation. (d) Bootstrap distribution of the mean absolute deviation of modularity from its mean, mean *|Q − µ|*, within each PGS group (half-violin: bootstrap density; box: median and interquartile range); modularity variability increases across the low, middle, and high PGS groups. (e) The same for global efficiency, mean *|E − µ|*, which shows no comparable broadening. (f) Joint distribution of modularity (Q) and global efficiency (E), both covariate-residualised, summarised per PGS group by a bootstrapped 95% covariance ellipse with its centroid and 95% confidence crosshairs; per-participant values appear as rug ticks on each axis. Network metrics were residualised for age, intracranial volume, and head motion.

This pattern showed clear network-level specificity. None of the continuous tests detected sex-modulated heteroscedasticity for global efficiency (variance-regression interaction *p* = 0.602; decile-trend difference *p* = 0.542; balanced-bootstrap trend difference *p* = 0.580), indicating that genetic predisposition specifically affects network segregation mechanisms whilst leaving integration properties more constrained. Together, these continuous analyses indicate that the broadening of viable modular configurations with increasing genetic predisposition is largely confined to male participants, whereas female network organisation appears comparatively stable across the full distribution of polygenic load.

### 3.2 No Systematic Shifts in Brain Organisation, Behaviour, or Their Association

Landscape theory predicts that genetic predisposition should expand variability without systematically shifting mean brain organisation or behaviour. Consistent with this prediction, polygenic scores showed no significant association with social-cognitive difficulties (*β* = 0.11, *p* = 0.209, *n* = 467; Figure 2c). Similarly, polygenic scores were not systematically associated with functional connectivity. Edge-wise analyses across the full sample (*N* = 910) revealed essentially no associations between polygenic scores and connectivity strength at any parcellation resolution after false discovery rate correction: no significant edges at the 50-node resolution (1,225 connections tested), a single edge at the 100-node resolution (4,950 connections), and no edges at the 200-node resolution (19,900 connections). Crucially, no polygenic-score × social-difficulty interactions survived correction at any resolution, indicating that genetic predisposition does not appear to moderate the edge-level relationship between connectivity and behaviour, or, at least, not strongly enough to be detectable with 900 participants. Social-cognitive difficulty itself, by contrast, was associated with a modest but consistent set of connectivity differences across resolutions (32, 30, and 18 significant edges at 50, 100, and 200 nodes, respectively, see Figure 3), confirming that the connectivity measures were sensitive enough to detect behaviourally relevant variation even when no polygenic-score signal was present. These edge-wise null findings for PGS, in contrast with the non-null findings for behaviour, are consistent with the landscape theory prediction that genetic predisposition influences the dispersion of neural configurations rather than their central tendency.

**Fig. 3.**
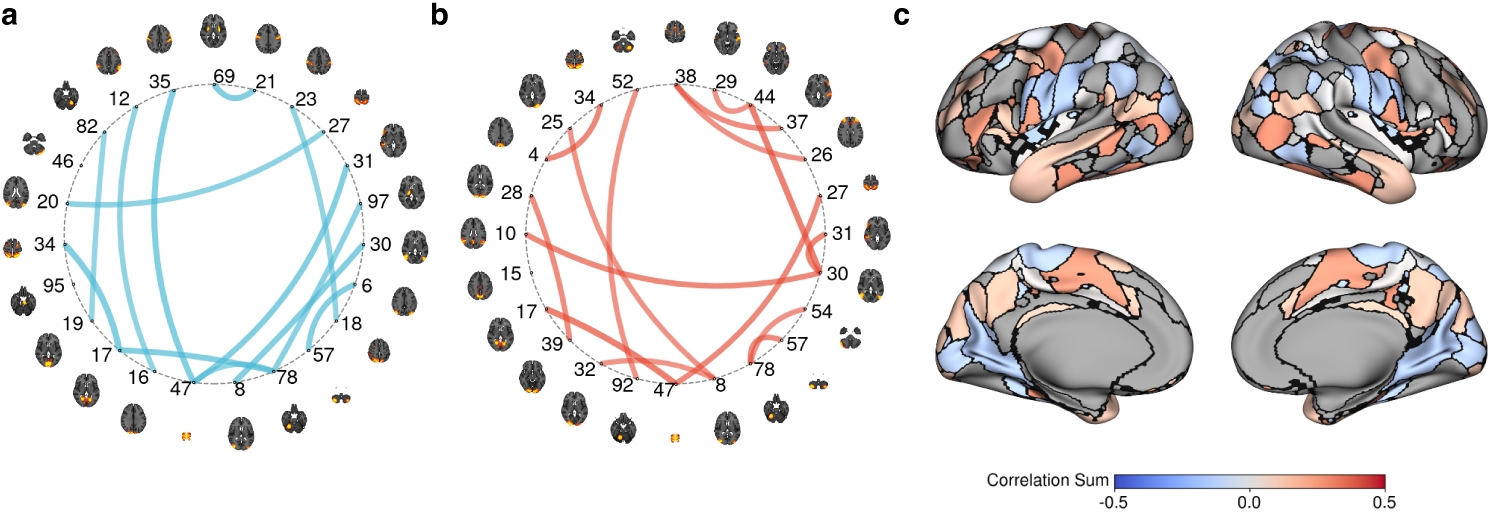
Univariate edge-level associations between resting-state functional connectivity and social difficulty. Each panel summarises the edge-wise relationship between functional connectivity (Fisher z-transformed; age, movement, intracranial volume, and sex regressed) and the social difficulty composite score across the analytic sample. (a) Edges showing a significant negative association, i.e. weaker connectivity in participants with higher social difficulty. (b) Edges showing a significant positive association, i.e. stronger connectivity in participants with higher social difficulty. (c) Per-node sum of suprathreshold edges, projected onto the cortical surface, indicating regions whose overall connectivity profile is most strongly implicated. In (a) and (b), nodes are placed at the peak MNI coordinates of the ICA components comprising the Human Connectome Project group parcellation and overlaid on axial template slices; edge colour denotes association sign and saturation scales with the magnitude of the standardised coefficient. All edges are thresholded at FDR q¡0.05

Mean network organisation itself did not predict social-cognitive functioning across the full sample. Regressing standardised PGS on each graph metric, sex, and their interaction revealed no evidence of association for either modularity (main effect *β* = −1.70, *p* = 0.239; sex × modularity *β* = 1.89, *p* = 0.362; *R*^2^ = 0.002) or global efficiency (main effect *β* = 1.09, *p* = 0.643; sex × global efficiency *β* = 0.28, *p* = 0.931; *R*^2^ = 0.001), with overall model fits indistinguishable from null (*F p* = 0.688 and 0.887, respectively). These convergent null findings on the means complement the variability results and are consistent with landscape theory’s prediction that genetic liability for autism is expressed through variability in network configuration rather than through shifts in average network organisation.

## 4 Discussion

Our findings provide partial support for landscape theories of neurodevelopmental diversity. Rather than uniformly shifting brain organisation toward dysfunction, autism genetic predisposition was associated with expanded variability in neural network configuration, though this expansion was confined to male participants. Across three convergent tests of continuous heteroscedasticity, male participants exhibited graded broadening of modular network configurations across the polygenic spectrum, whereas female participants showed no comparable expansion. Neither sex showed systematic differences in mean network organisation or in mean associations with social-cognitive functioning. This pattern of increased dispersion without mean shifts, observed in male participants, is consistent with what landscape theory predicts when genetic factors loosen developmental constraints rather than deterministically altering neural architecture [3, 4]; the absence of comparable expansion in female participants, however, requires that this framework be qualified by considerations of sex-differential canalisation, sex-specific polygenic score calibration [32], or both.

The specificity of variance effects to modularity, rather than global efficiency, suggests that genetic predisposition particularly influences the brain’s capacity to form segregated functional communities, at least in male participants. Network segregation supports specialised processing within distinct neural systems, whereas integration facilitates communication between them [33]. Our finding that modularity variance expands while efficiency variance remains stable implies that genetic predisposition affects the balance between these organisational principles in heterogeneous ways across male individuals. This dissociation emerged consistently across multiple independent tests of continuous heteroscedasticity spanning the full polygenic distribution, indicating that the variance-modularity relationship is a graded property of the genetic spectrum in males. Among male participants, some individuals with high genetic predisposition may develop highly segregated networks, others more integrated configurations, while those with low genetic predisposition converge toward a narrower range of solutions that balance segregation and integration. This interpretation aligns with the Anna Karenina principle articulated by Finn and colleagues: optimal configurations are constrained, but suboptimal ones can deviate in multiple directions [4]. Whether this principle operates symmetrically across sexes or, as our results suggest, primarily in males, is a question to which we return below.

The absence of systematic brain-behaviour associations at the sample level is itself theoretically informative. Neither polygenic score nor mean network organisation predicted social-cognitive functioning in our sample, and these null findings despite the robust case-control prediction achieved in clinical GWAS [5] are precisely what landscape theory anticipates. If genetic predisposition expands the range of viable neural configurations rather than shifting their central tendency, aggregate scores should poorly predict individual outcomes, and mean-level network measures should likewise show weak behavioural correspondence. Within this framework, phenotypic heterogeneity reflects meaningful biological variability in the trajectories that translate genetic predisposition into neural organisation: two individuals with comparable polygenic scores may develop markedly different network configurations, with behavioural outcomes depending on the match between achieved configuration and task demands rather than on genetic load per se. Notably, the sex-dependent broadening of modular configurations in males did not translate into a detectable sex-specific brain–behaviour relationship in this sample. Whether male-specific configurational diversity confers particular behavioural advantages or challenges is a question that will likely require substantially larger samples or analytical approaches tailored to non-linear, subgroup-specific patterns.

Why might landscape-broadening effects of polygenic load manifest in males but not in females? Two non-mutually-exclusive interpretations warrant consideration. The first is biological: the female protective effect proposes that females are buffered against the phenotypic expression of autism-associated genetic load, requiring greater cumulative liability before clinical features emerge [34, 35]. If this protection extends to neural organisation itself, e.g. through stronger developmental canalisation, sex-differential gene expression, or hormonal modulation of synaptic development, then the same polygenic load that expands the landscape of viable male network configurations may be absorbed in females without producing comparable organisational diversification. The female protective effect would thus operate at the level of network organisation as well as at the level of clinical diagnosis, with female neural development exhibiting a narrower distribution of viable configurations across the polygenic spectrum. The second interpretation is methodological: the GWAS underlying our polygenic scores [5] was derived from a clinical sample with a substantial male representation, reflecting the well-documented sex bias in autism diagnosis, particularly in cohorts that were diagnosed before the female presentation of autism was better understood and widely recognised. The variants identified, and consequently the polygenic scores constructed from them, may be more strongly calibrated to autism-relevant biology in males than in females, in whom the genetic architecture of the autism phenotype appears to differ in important respects [36, 37]. Under this interpretation, the female null reflects not biological resilience but a measurement asymmetry inherent to current PGS construction. Indeed, the genetic correlation between male- and female-stratified autism GWAS is high but significantly below 1 (rg = 0.80, SE = 0.08, [32]). However, distinguishing these accounts will require sex-stratified GWAS of sufficient power to construct sex-specific polygenic scores, an undertaking that remains beyond the scope of currently available autism genomic resources but represents a critical next step for landscape-theory research.

Several mechanisms could underlie the increased neural variability observed in male individuals with high genetic predisposition. Developmental compensation represents one possibility, whereby alternative neural pathways emerge when typical circuits are disrupted [38]. Neuroimaging studies in autism have documented compensatory recruitment patterns, including increased right-hemisphere activation during working memory [39], enhanced hippocampal engagement during memory tasks [40], and recruitment of additional regions during language processing [41]. Such compensation could manifest as reduced modularity if regions typically confined to distinct modules become integrated into broader processing networks. Alternatively, increased variability may reflect developmental stochasticity rather than active compensation. Genetic factors could introduce noise into developmental processes, leading to more variable outcomes without implying functional optimisation. A third possibility is that genetic predisposition opens access to qualitatively different but equally viable neural solutions, consistent with Johnson’s interactive specialisation framework [42]. These mechanisms are not mutually exclusive; the expanded developmental landscape may accommodate compensated trajectories, stochastic deviations, and alternative solutions simultaneously. That such broadening was not detectable in female participants implies that whichever mechanisms operate, they are expressed differently, or are more tightly constrained, across sexes.

Our findings speak to ongoing reconceptualisations of neurodevelopmental conditions within a neurodiversity framework. The observation that genetic predisposition expands neural variability in males rather than uniformly producing dysfunction challenges deficit-focused models that frame autism primarily in terms of impairment [43]. Instead, our results suggest that autism-associated genetic variants contribute to a broader repertoire of neural configurations in male non-autistic adults, some of which may confer advantages in specific contexts while others may present challenges. This perspective does not minimise the genuine difficulties many autistic individuals experience, but it does reframe phenotypic heterogeneity as reflecting meaningful biological diversity rather than measurement error or incomplete genetic models. The substantial overlap in neural organisation between individuals with high and low genetic predisposition, observed in both sexes, underscores that genetic load alone does not determine neural or behavioural outcomes.

These results have methodological implications for autism neuroimaging research. The increased neural idiosyncrasy documented in clinical samples [8–12] has typically been interpreted as reflecting autism-specific neurobiology, but our findings suggest that some portion of this variability, at least among male participants, may be attributable to genetic predisposition present across the population continuum. Case-control designs that compare autistic individuals against non-autistic controls confound diagnostic status with genetic load, potentially attributing to autism per se what is more accurately a consequence of elevated genetic predisposition. The sex-dependence of this association adds a further consideration: aggregating across sexes in case-control comparisons may obscure relationships that operate primarily in one sex, an issue compounded by the heavily male-skewed composition of most clinical autism samples. Future studies may benefit from incorporating polygenic scores, stratifying analyses by sex, or both, to partition variance attributable to common genetic factors from that arising through other mechanisms, including rare variants, environmental factors, and gene-environment interactions [44].

Several limitations should be acknowledged. First, the cross-sectional design cannot establish whether the increased neural variability observed in high-PGS male individuals reflects different developmental trajectories, compensatory processes, or stable individual differences present from early life. Longitudinal studies following individuals from infancy would better illuminate how genetic predisposition shapes the emergence of neural organisation over time, and whether the sex-asymmetric pattern observed here is present from the outset or emerges during pubertal or post-pubertal development. Second, the effect sizes observed here are modest for both the male-specific variance broadening and for the null mean-level associations, which is characteristic of imaging-genetics research more broadly but precludes meaningful clinical translation at the individual level. Third, the male-specific variance broadening, while convergent across three independent analytical strategies, requires replication in independent samples before strong conclusions are warranted. Fourth, the female null result must be interpreted with appropriate caution: with approximately half of the analytic sample available for each sex, statistical power to detect a sex-specific expansion is reduced relative to the full-sample analyses, and stable estimation of variance changes across the polygenic distribution requires substantial sample sizes. The sex × PGS interaction itself is the more reliable inference; a true but smaller female effect, or an effect operating in a different direction, cannot be excluded on the basis of these data alone. Fifth, examining whether genetic predisposition moderates brain–behaviour relationships, as landscape theory would predict, requires substantially larger samples than currently available in imaging-genetics datasets, particularly when sex stratification is also required. Sixth, our polygenic scores derive from a genome-wide association study conducted predominantly in individuals of European ancestry [5], limiting generalisability across populations with different genetic architectures. Future work should examine whether landscape theory predictions hold across diverse genetic backgrounds. Seventh, our sample excluded individuals with psychiatric or neurological diagnoses, meaning that our high-PGS group may have included individuals with subclinical autistic traits or the broader autism phenotype but not those who would meet diagnostic criteria. The neural variability we observe may therefore represent the upper bound of successful adaptation to genetic predisposition in non-clinical populations. Whether similar patterns of expanded variability characterise autistic individuals, and whether the male-specificity observed here generalises to or differs in clinical populations, remains an important question for future research. Eighth, our genetic analysis focused on polygenic scores derived from common single-nucleotide polymorphisms, capturing only one component of the genetic architecture of autism. Rare variants with high penetrance, including de novo copy number variants and loss-of-function mutations, have been strongly implicated in autism aetiology [44] and may exert distinct effects on neural organisation that are not reflected in common-variant polygenic scores. Whether landscape theory predictions regarding expanded neural variability generalise to individuals whose genetic predisposition is driven primarily by rare, high-impact variants remains an open question. Finally, our social-cognitive measure was based on reaction times from experimental tasks rather than real-world social functioning or autism-specific assessments. While this approach captures processing efficiency, it may not reflect the social challenges most relevant to autistic individuals’ daily lives.

## 5 Conclusion

This study demonstrates that autism genetic predisposition is associated with expanded variability in brain network organisation rather than systematic shifts toward dysfunction. Within the male subsample, individuals with high polygenic scores showed substantially increased variance in network modularity than those with low genetic load, while mean network organisation and social-cognitive functioning did not differ between groups. These findings provide direct empirical support for landscape theories of neurodevelopmental diversity, suggesting that genetic predisposition loosens constraints on neural development, enabling a broader range of configurations – some optimal, others less so. That neither polygenic score nor mean network organisation predicted social-cognitive performance at the sample level is consistent with this interpretation: where genetic effects operate on the dispersion rather than the central tendency of neural configurations, behavioural relevance is unlikely to emerge from aggregate associations and should instead depend on individual developmental trajectories. This work reframes phenotypic heterogeneity in autism as reflecting meaningful biological diversity in how genetic predisposition translates into neural organisation, with implications for understanding neurodevelopmental variation across the population spectrum.

## Acknowledgements

The authors wish to thank the Human Connectome Project team and all research participants for providing the rich and detailed neuroimaging, behavioural, and genetic data that made this study possible. We are also grateful to the iPSYCH study team for making the polygenic score data publicly available. Finally, we thank Tinka Beemsterboer, data steward at the University of Amsterdam, for her support in designing the data management plan for this study.

## Declarations

The authors declare no conflict of interest, financial or otherwise.

## Code availability

All analysis code used in this study is publicly available on GitHub (https://github.com/joebathelt/AutismBrainVariability). The repository contains the analysis code and associated configuration files that reproduce all preprocessing, statistical, and visualisation procedures described in the Methods section. The repository contains an environment file that includes all software dependencies and package versions, allowing for exact replication of the analytical pipeline from raw data processing through final figure generation.

## Data availability

The neuroimaging, behavioural, and demographic data from the Human Connectome Project Young Adult sample (HCP-YA) used in this study are publicly available through the HCP website (https://www.humanconnectome.org), subject to completion of a data use agreement. However, genetic data from the HCP-YA, including the genotyping array data used to calculate polygenic scores in the present analyses, are available under controlled access provisions that require submission of a formal data access application to the HCP consortium. In accordance with HCP data usage requirements, all participant IDs in the current study have been replaced with study-specific IDs. For the exact computational replication of the current results, a file that matches our study-specific IDs to the HCP-YA IDs has been submitted to the HCP consortium. Summary statistics for the autism genome-wide association study (GWAS) used to construct the polygenic scores were obtained from the iPsych consortium and are publicly available through the PGS catalogue website (https://www.pgscatalog.org/score/PGS000327/).

## Author contribution

**JB**: Conceptualization, Formal analysis, Methodology, Project administration, Software, Supervision, Visualization, Writing – original draft, Writing – review & editing. **DM**: Conceptualization, Formal analysis, Methodology, Software, Writing – original draft, Writing – review & editing. **HMG**: Supervision, Writing – review & editing

## Appendix A Data inclusion

See Figure S1 for an overview of data inclusion.

### Calculation of the social processing factor - Detailed Methods

The HCP-YA protocol included broad assessment of behaviour and cognition, but did not contain any autism-specific assessments. We chose to focus specifically on social processing because difficulties in social communication and interaction constitute the defining diagnostic feature of autism [45], and are most directly relevant to the phenotype under investigation. Although autism is also associated with differences in executive function and language [46, 47], including these broader cognitive domains risked producing a general factor of cognitive ability rather than one that captures variation specifically relevant to the autism phenotype. Therefore, we developed a factor score of social processing based on existing measurements. We employed a four-step process: first, selecting cognitive and behavioural measures based on theoretical understanding; second, processing the data to handle missing values and extreme data points, and create a training and test split; three, identifying measures that formed a coherent social processing factor through exploratory factor analysis; and finally, evaluating the winning model’s fit using confirmatory factor analysis. The following sections describe each step in more detail.

**Fig. S1.**
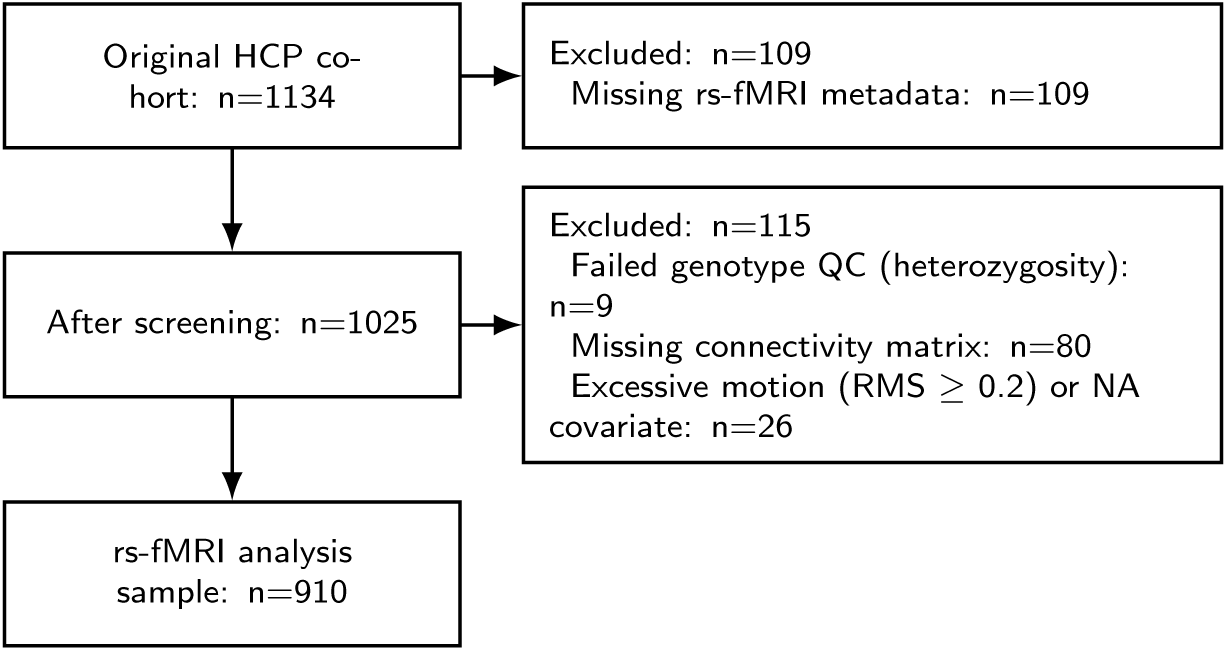
Participant flow diagram showing sample sizes for different analyses. PGS = polygenic score; SD = Standard Deviation; rs-fMRI = resting-state functional magnetic resonance imaging; QC: quality control.

#### A.1 Selecting cognitive and behavioural measures

The behavioural protocol of HCP-YA sample included measures of the NIH Toolbox, custom questionnaires, and behavioural scores from task fMRI assessments. The full protocol with a description of all measures is available from the HCP website (https://humanconnectome.org). Our selection criteria were that measures had to be numeric and be related to social, emotional, or language processing based on the descriptions of NIH Toolbox documentation or the HCP task-based fMRI paper [48].

##### Penn Emotion Recognition 40

The Penn Emotion Recognition Task (ER40) is a computerized neuropsychological test designed to assess facial emotion recognition abilities [49]. It includes 40 colour photographs of adult faces (varying in race and sex), each showing one of four basic emotions (sad, happy, angry, or scared) or a neutral expression, with eight faces per expression. Participants were presented with a face and had to choose a label. Accuracy and reaction time were recorded. Facial emotion recognition is consistently reported to be different in autistic individuals, with meta-analytic evidence indicating moderate deficits across multiple emotional expressions [50, 51].

##### Language processing

For this fMRI task, participants heard short auditory stories (5–9 sentences) based on Aesop’s fables, followed by a two-alternative forced-choice question about the story’s topic. For instance, after a story about an eagle that saves a man who had done him a favor, participants were asked, “That was about revenge or reciprocity?” [48]. Response accuracy and reaction time were recorded. We included this task because the questions required reasoning about the intentions of characters, a capacity closely linked to theory of mind, a domain in which autistic individuals often show performance differences [52].

##### Emotion processing

During this fMRI task, participants viewed photographs of angry or fearful facial expressions and were asked to choose which of the two images at the bottom matched the one shown at the top. We included this task because it taps face processing abilities that may be different in autistic individuals [50, 51].

##### Social processing

While in the fMRI scanner, participants watched short video clips (20 seconds) showing shapes (squares, circles, and triangles) either interacting meaningfully or moving randomly. The clips were followed by a question asking participants to choose one of three options: whether the shapes were socially interacting, whether they were unsure, or whether there was no interaction. Response accuracy and reaction time were recorded. This task draws on the classic [53] paradigm, in which observers spontaneously attribute mental states to geometric shapes based on their movement patterns. Autistic individuals have been shown to produce fewer mental state attributions when viewing such animations [54, 55].

##### NIH Toolbox Social Relationships

Participants completed the NIH Toolbox questionnaire on social relationships, which included items in 6 domains assessing their views on friendships, feelings of loneliness, perceived hostility and rejection from others, as well as the emotional and instrumental support they received. Individual items in each scale were summed to create a domain score. These domains are highly relevant to autism, as autistic individuals report higher levels of loneliness and lower friendship quality compared to non-autistic peers [56, 57].

#### A.2 Data processing

Data processing involved handling missing values and extreme data points, followed by transformation and scaling. We removed participants with extreme missingness (50% missing across >50% of features), reducing overall missing value rates (max. 5.6%). The remaining data were scaled, and outliers (outside 1.5 × interquartile range) were marked as missing. Missing value imputation was performed using the ‘ImputeKernel’ function from the ‘miceforest’ package v6.0.1 in Python, which builds gradient boosting models to iteratively predict missing values from other data. The results were pooled over 5 iterations to account for imputation uncertainty. Following these preprocessing steps, the sample was randomly split into a training set (80%) and a test set (20%).

#### A.3 Creating a social processing factor

To identify the latent structure underlying social processing, we performed an exploratory factor analysis (EFA) on the training data, specifying a single-factor solution. We evaluated model fit using the Chi-square test (*p >* 0.05 for good fit), the Root Mean Square Error of Approximation (RMSEA; ≤ 0.05 for good fit), the Tucker–Lewis Index (TLI; values near 0.9 for good fit), and the Root Mean Square Residual (RMSR; < 0.08 desirable), which served as relative indices of approximate fit. To compare across models, we used the Bayesian Information Criterion (BIC), with lower values indicating better relative fit.

#### A.4 Evaluation of the social processing factor

We conducted confirmatory factor analysis (CFA) on the test set to evaluate the model identified through exploratory factor analysis in the training set. Model fit was assessed using the same approximate fit indices described above, where applicable, along with the Comparative Fit Index (CFI; values close to 0.90 or higher for good fit and should exceed the TLI), and the Standardised Root Mean Square Residual (SRMR; < 0.08 for good fit). For model comparison, we used the Akaike Information Criterion (AIC), with lower values indicating better relative fit, in addition to BIC. Factor loadings were tested for statistical significance, with one loading fixed at 1.0 to anchor the scale and aid interpretation. Finally, the CFA model was fit to the full dataset to derive social processing scores for the entire sample.

### Supplementary Methods: Polygenic Score Pipeline

#### A.5 Software and dependencies

The genetics pipeline was implemented with the following software versions: PLINK 2.0 [20], `GCTA v1.94.3` [19], `PRSice-2 v2.3.5` [18], and R (with `plinkQC` [17], `ggplot2`, and `data.table`). Downstream evaluation scripts were written in Python with `pandas`, `numpy`, `scipy`, `scikit-learn`, `statsmodels`, `seaborn`, `matplotlib`, and `pingouin`. The full analysis pipeline was executed inside a conda environment and integrated using Snakemake. The full analysis code and environment configuration files are available via the project repository.

#### A.6 Target-data genotype quality control

Target-data quality control was performed with `plinkQC` [17], which implements the standard per-individual and per-marker pipeline established for GWAS and PGS applications. Per-individual filters comprised: sample missingness (< 3%), sex verification (agreement between the sex recorded in the HCP demographic data and the sex coded in the genotype files, with discordant individuals removed; the chromosomal-sex F-statistic was *not* used for exclusion, as the HCP genotype sex codes proved unreliable—see Results), and heterozygosity outliers (beyond ±3 SD of the sample mean, conditional on sample missingness). Per-marker filters comprised: SNP missingness (< 1%), Hardy–Weinberg equilibrium (*p >* 10*^−^*^6^), and MAF (≥ 0.01). An initial relatedness check within `plinkQC` flagged individuals with *π̂* > 0.1875 using IBD statistics computed on LD-pruned SNPs; relatedness-based exclusion of individuals, however, was not applied at the QC stage but handled separately during unrelated-subset calibration (see Supplementary Methods S5). High-LD genomic regions were masked during IBD estimation, in line with `plinkQC` defaults for genome build hg19.

#### A.7 Base-data processing and SNP harmonisation

Base-data preparation of the iPSYCH autism GWAS summary statistics [5] consisted of three sequential steps: removal of duplicate SNPs (retaining the first occurrence of each variant identifier), exclusion of SNPs with missing effect-size or standard-error estimates or imputation quality < 0.7, and exclusion of SNPs with base-data MAF < 0.01. Harmonisation of target and base data was performed in R using a custom script that compared allele codes across datasets on a per-SNP basis. Variants with unambiguous strand matches were retained with their base-data effect allele as the PLINK A1 reference; variants with strand-flip-compatible allele codes were recoded to match the base-data strand; palindromic (A/T and C/G) SNPs and variants with irreconcilable allele codes were written to an exclusion list and removed from the final PGS SNP set using PLINK’s `--exclude` flag. Heterozygosity statistics computed on the pruned target-data SNP set were used during harmonisation to verify that allele recoding did not introduce systematic bias in genotype frequencies.

#### A.8 PGS threshold selection and unrelated-subset calibration

PGS were computed across the default set of PRSice-2 p-value thresholds using the `--no-regress` flag, which writes scores at all thresholds (`.all score` output) rather than performing target-data regression-based selection. The target threshold (*p* = 0.1) was pre-specified on the basis of the discovery GWAS [5] and fixed prior to downstream analyses. Clumping parameters followed PRSice-2 defaults (*p*_1_ = 1, *p*_2_ = 1, clump window = 250 kb, clump *r*^2^ = 0.1). The unrelated subset was defined conjunctively through: (i) phenotypic status as a non-twin in the HCP twin structure; (ii) a PLINK`--rel-cutoff` of 0.125 applied to the LD-pruned target SNP set; and (iii) availability of complete social-processing factor scores.

#### A.9 BLUP extension to the full sample

Extension of PGS to the full sample including related individuals was implemented in GCTA v1.94.3 [19] via the following sequence of calls. A genomic relationship matrix was computed on autosomal markers only (`gcta64 --bfile <target> --autosome --make-grm`). A linear mixed model was fitted to the standardised unrelated-sample PGS, treating the GRM as the random-effect design matrix (`--grm <grm> --pheno <unrelated pgs> --reml --reml-pred-rand`), which yielded individual-level random-effect predictions (`.indi.blp`) for all individuals in the GRM, including those excluded from the calibration subset. Per-SNP BLUP effect sizes were then back-calculated from the individual predictions (`--blup-snp <indi.blp> --autosome`), producing an effect-size file (`.snp.blp`) compatible with PLINK scoring. Finally, the full-sample BLUP-extended PGS was computed by PLINK with the `–score <snp.blp> sum` flag, which sums signed per-SNP contributions across all autosomal markers and writes one score per individual to the `.profile` output. This procedure inherits the random-effects framework from the LMM fit and therefore produces scores that are calibrated on the unrelated sample while accommodating the familial structure of the full HCP cohort.

#### A.10 Residualisation and Standardisation

For association and group-level analyses, the BLUP-extended PGS was residualised with respect to age and the first 5 principal components in an ordinary-least-squares regression fitted to the full sample. Residuals were standardised (*z*-scored).

## References

[1] Taylor MJ, Martin J, Lu Y, Brikell I, Lundström S, Larsson H, et al. Association of genetic risk factors for psychiatric disorders and traits of these disorders in a Swedish population twin sample. JAMA Psychiatry. 2019 Mar;76(3):280–289.

[2] Schendel D, Munk Laursen T, Albiñana C, Vilhjalmsson B, Ladd-Acosta C, Fallin MD, et al. Evaluating the interrelations between the autism polygenic score and psychiatric family history in risk for autism. Autism Res. 2022 Jan;15(1):171–182.

[3] van den Heuvel MP, Sporns O. A cross-disorder connectome landscape of brain dysconnectivity. Nat Rev Neurosci. 2019;20(7):435–446.

[4] Finn ES, Glerean E, Khojandi AY, Nielson D, Molfese PJ, Handwerker DA, et al. Idiosynchrony: From shared responses to individual differences during naturalistic neuroimaging. Neuroimage. 2020;215:116828.

[5] Grove J, Ripke S, Als TD, Mattheisen M, Walters RK, Won H, et al. Identification of common genetic risk variants for autism spectrum disorder. Nat Genet. 2019 Mar;51(3):431–444.

[6] de Wit MM, Morgan MJ, Libedinsky I, Austerberry C, Begeer S, Abdellaoui A, et al. Systematic review and meta-analysis: Phenotypic correlates of the autism polygenic score. JAACAP Open. 2025 Apr;.

[7] Di Martino A, Castellanos FX, Anderson J, others. The Autism Brain Imaging Data Exchange (ABIDE) consortium: open sharing of autism resting state fMRI data. Brain Mapping, Date 2012;.

[8] Benkarim O, Paquola C, Park BY, Hong SJ, Royer J, Vos de Wael R, et al. Connectivity alterations in autism reflect functional idiosyncrasy. Commun Biol. 2021 Sep;4(1):1078.

[9] Dickie EW, Ameis SH, Shahab S, Calarco N, Smith DE, Miranda D, et al. Personalized Intrinsic Network Topography mapping and functional connectivity deficits in autism spectrum disorder. Biol Psychiatry. 2018 Aug;84(4):278–286.

[10] Xie Y, Sun J, Man W, Zhang Z, Zhang N. Personalized estimates of brain cortical structural variability in individuals with Autism spectrum disorder: the predictor of brain age and neurobiology relevance. Mol Autism. 2023 Jul;14(1):27.

[11] Ecker C, Pretzsch CM, Bletsch A, Mann C, Schaefer T, Ambrosino S, et al. Interindividual differences in cortical thickness and their genomic underpinnings in autism spectrum disorder. Am J Psychiatry. 2022 Mar;179(3):242–254.

[12] Tung YH, Lin HY, Chen CL, Shang CY, Yang LY, Hsu YC, et al. Whole brain white matter tract deviation and idiosyncrasy from normative development in autism and ADHD and unaffected siblings link with dimensions of psychopathology and cognition. Am J Psychiatry. 2021 Aug;178(8):730–743.

[13] Mihailov A, Philippe C, Gloaguen A, Grigis A, Laidi C, Piguet C, et al. Cortical signatures in behaviorally clustered autistic traits subgroups: a population-based study. Transl Psychiatry. 2020 Jun;10(1):207.

[14] Khachadourian V, Mahjani B, Sandin S, Kolevzon A, Buxbaum JD, Reichenberg A, et al. Comorbidities in autism spectrum disorder and their etiologies. Transl Psychiatry. 2023 Feb;13(1):71.

[15] Turiaco F, Arnone F, Drago A, Muscatello MRA, Bruno A, Iannuzzo F. Escitalopram and functional connectivity in major depressive disorder: A systematic review. J Psychopharmacol. 2025 Oct;39(10):1062–1071.

[16] Van Essen DC, Ugurbil K, Auerbach E, Barch D, Behrens TEJ, Bucholz R, et al. The Human Connectome Project: a data acquisition perspective. Neuroimage. 2012 Oct;62(4):2222–2231.

[17] Syed M, Walter C, Meyer HV. PlinkQC: An integrated tool for ancestry inference, sample selection, and quality control in population genetics. bioRxivorg. 2025 Nov;.

[18] Choi SW, O’Reilly P. PRSice 2: POLYGENIC RISK SCORE SOFTWARE (UPDATED) AND ITS APPLICATION TO CROSS-TRAIT ANALYSES. Eur Neuropsychopharmacol. 2019 Jan;29:S832.

[19] Yang J, Lee SH, Goddard ME, Visscher PM. GCTA: a tool for genome-wide complex trait analysis. Am J Hum Genet. 2011 Jan;88(1):76–82.

[20] Chang CC, Chow CC, Tellier LC, Vattikuti S, Purcell SM, Lee JJ. Second-generation PLINK: rising to the challenge of larger and richer datasets. Gigascience. 2015;4(1):s13742–015.

[21] Smith SM, Beckmann CF, Andersson J, Auerbach EJ, Bijsterbosch J, Douaud G, et al. Resting-state fMRI in the Human Connectome Project. Neuroimage. 2013 Oct;80:144–168.

[22] Glasser MF, Sotiropoulos SN, Wilson JA, Coalson TS, Fischl B, Andersson JL, et al. The minimal preprocessing pipelines for the Human Connectome Project. Neuroimage. 2013 Oct;80(C):105 124.

[23] Filippini N, MacIntosh BJ, Hough MG, Goodwin GM, Frisoni GB, Ebmeier K, et al. Distinct patterns of brain activity in young carriers of the APOE e4 allele. Neuroimage. 2009 Jul;47:S139.

[24] Rubinov M, Sporns O. Complex network measures of brain connectivity: Uses and interpretations. Neuroimage. 2010 Sep;52(3):1059 1069.

[25] Garrison KA, Scheinost D, Finn ES, Shen X, Constable RT. The (in)stability of functional brain network measures across thresholds. Neuroimage. 2015 Sep;118:651–661.

[26] Smith SM, Nichols TE, Vidaurre D, Winkler AM, Behrens TEJ, Glasser MF, et al. A positive-negative mode of population covariation links brain connectivity, demographics and behavior. Nat Neurosci. 2015 Nov;18(11):1565–1567.

[27] Yeo BTT, Krienen FM, Sepulcre J, Sabuncu MR, Lashkari D, Hollinshead MO, et al. The organization of the human cerebral cortex estimated by intrinsic functional connectivity. Journal of Neurophysiology. 2011;.

[28] Assaf M, Jagannathan K, Calhoun VD, Miller L, Stevens MC, Sahl R, et al. Abnormal functional connectivity of default mode sub-networks in autism spectrum disorder patients. Neuroimage. 2010 Oct;53(1):247–256.

[29] Bathelt J, Geurts HM. Difference in default mode network subsystems in autism across childhood and adolescence. Autism. 2021 Feb;25(2):556–565.

[30] Chen YY, Uljarevic M, Neal J, Greening S, Yim H, Lee TH. Excessive functional coupling with less variability between salience and default mode networks in autism spectrum disorder. Biol Psychiatry Cogn Neurosci Neuroimaging. 2022 Sep;7(9):876–884.

[31] Bullmore E, Sporns O. The economy of brain network organization. Nat Rev Neurosci. 2012 Apr;13(5):336–349.

[32] Gu Y, Maria-Stauffer E, Bedford SA, APEX consortium, iPSYCH-autism consortium, Romero-Garcia R, et al. Polygenic scores for autism are associated with reduced neurite density in adults and children from the general population. Mol Psychiatry. 2025 Aug;30(8):3393–3403.

[33] Sporns O. Network attributes for segregation and integration in the human brain. Curr Opin Neurobiol. 2013 Apr;23(2):162 171.

[34] Robinson EB, Lichtenstein P, Anckarsäter H, Happé F, Ronald A. Examining and interpreting the female protective effect against autistic behavior. Proc Natl Acad Sci U S A. 2013 Mar;110(13):5258–5262.

[35] Werling DM. The role of sex-differential biology in risk for autism spectrum disorder. Biol Sex Differ. 2016 Nov;7(1):58.

[36] Mitra I, Tsang K, Ladd-Acosta C, Croen LA, Aldinger KA, Hendren RL, et al. Pleiotropic mechanisms indicated for sex differences in autism. PLoS Genet. 2016 Nov;12(11):e1006425.

[37] Antaki D, Guevara J, Maihofer AX, Klein M, Gujral M, Grove J, et al. A phenotypic spectrum of autism is attributable to the combined effects of rare variants, polygenic risk and sex. Nature genetics. 2022;54(9):1284–1292.

[38] Livingston LA, Colvert E, Social Relationships Study Team, Bolton P, Happé F. Good social skills despite poor theory of mind: exploring compensation in autism spectrum disorder. J Child Psychol Psychiatry. 2019 Jan;60(1):102–110.

[39] Yeung MK, Lee TL, Chan AS. Right-lateralized frontal activation underlies successful updating of verbal working memory in adolescents with high-functioning autism spectrum disorder. Biol Psychol. 2019 Nov;148(107743):107743.

[40] Hogeveen J, Krug MK, Geddert RM, Ragland JD, Solomon M. Compensatory hippocampal recruitment supports preserved episodic memory in autism spectrum disorder. Biol Psychiatry Cogn Neurosci Neuroimaging. 2020 Jan;5(1):97– 109.

[41] Eigsti IM, Stevens MC, Schultz RT, Barton M, Kelley E, Naigles L, et al. Language comprehension and brain function in individuals with an optimal outcome from autism. NeuroImage Clin. 2016;10:182–191.

[42] Johnson MH. Autism as an adaptive common variant pathway for human brain development. Dev Cogn Neurosci. 2017;25:5–11.

[43] Fletcher-Watson S. Transdiagnostic research and the neurodiversity paradigm: commentary on the transdiagnostic revolution in neurodevelopmental disorders by Astle et al. J Child Psychol Psychiatry. 2022 Apr;63(4):418–420.

[44] Bourgeron T. Current knowledge on the genetics of autism and propositions for future research. C R Biol. 2016 Jul;339(7-8):300–307.

[45] American Psychiatric Association. Diagnostic and Statistical Manual of Mental Disorders: DSM-5-TR. 5th ed. Washington, DC: American Psychiatric Publishing; 2022.

[46] Demetriou EA, Lampit A, Quintana DS, Naismith SL, Song YJC, Pye JE, et al. Autism spectrum disorders: a meta-analysis of executive function. Mol Psychiatry. 2018 May;23(5):1198–1204.

[47] Kwok EYL, Brown HM, Smyth RE, Oram Cardy J. Meta-analysis of receptive and expressive language skills in autism spectrum disorder. Res Autism Spectr Disord. 2015 Jan;9:202–222.

[48] Barch DM, Burgess GC, Harms MP, Petersen SE, Schlaggar BL, Corbetta M, et al. Function in the human connectome: task-fMRI and individual differences in behavior. Neuroimage. 2013 Oct;80:169–189.

[49] Gur RC, Ragland JD, Moberg PJ, Turner TH, Bilker WB, Kohler C, et al. Computerized neurocognitive scanning: I. Methodology and validation in healthy people. Neuropsychopharmacology. 2001 Nov;25(5):766–776.

[50] Uljarevic M, Hamilton A. Recognition of emotions in autism: a formal meta-analysis. J Autism Dev Disord. 2013 Jul;43(7):1517–1526.

[51] Lozier LM, Vanmeter JW, Marsh AA. Impairments in facial affect recognition associated with autism spectrum disorders: a meta-analysis. Dev Psychopathol. 2014 Nov;26(4 Pt 1):933–945.

[52] Gao S, Wang X, Su Y. Examining whether adults with autism spectrum disorder encounter multiple problems in theory of mind: a study based on meta-analysis. Psychon Bull Rev. 2023 Oct;30(5):1740–1758.

[53] Heider F. Social perception and phenomenal causality. Psychol Rev. 1944 Nov;51(6):358–374.

[54] Rasmussen CE, Jiang YV. Judging social interaction in the Heider and Simmel movie. Q J Exp Psychol (Hove). 2019 Sep;72(9):2350–2361.

[55] Castelli F. Autism, Asperger syndrome and brain mechanisms for the attribution of mental states to animated shapes. Brain. 2002 Aug;125(8):1839–1849.

[56] Mazurek MO. Loneliness, friendship, and well-being in adults with autism spectrum disorders. Autism. 2014;18(3):223–232.

[57] White SW, Roberson-Nay R. Anxiety, Social Deficits, and Loneliness in Youth with Autism Spectrum Disorders. J Autism Dev Disord. 2009;39(7):1006–1013.

